# Rubisco is slow across the tree of life

**DOI:** 10.1101/2025.01.19.633714

**Authors:** Benoit de Pins, Cyril Malbranke, Jagoda Jabłońska, Assaf Shmuel, Itai Sharon, Anne-Florence Bitbol, Oliver Mueller-Cajar, Elad Noor, Ron Milo

**Author notes:** Currently: Department of Biology, University of Naples Federico II, Naples 80126, Italy.

## Abstract

Rubisco is the main gateway through which inorganic carbon enters the biosphere, catalyzing the vast majority of carbon fixation on Earth. This pivotal enzyme has long been observed to be kinetically constrained. Yet, this impression is based on kinetic measurements heavily focused on eukaryotic rubiscos, a rather conserved group of low genetic diversity. Moreover, the fastest rubiscos that we know of so far were found among the sparsely sampled prokaryotes. Could there be yet faster rubiscos among the uncharted regions of rubisco’s phylogenetic diversity? Here, we perform a characterization of more than 250 rubiscos from a wide range of bacteria and archaea, thereby doubling the coverage of the diversity of this key enzyme. We assess the distribution of the carboxylation rates at saturating levels of CO_2_, and establish that rubisco is a relatively slow enzyme across the tree of life, never exceeding ≈30 reactions per second at 30°C. We show that relatively faster subclades share similar evolutionary contexts, involving micro-oxygenic environments or a CO_2_ concentrating mechanism. Leveraging a simple machine learning model trained on this dataset, we predict the carboxylation rate for all ≈68,000 sequenced rubisco variants found in nature to date. This study provides the largest and most diverse dataset of natural variants for an enzyme and their associated rates, establishing a solid benchmark for future efforts to predict catalytic rates from sequence data.

**Significance:** Discovering a fast carboxylating rubisco has been a long-standing challenge in the scientific community, given its potential impact on sustainable food and fuel production. Yet, only a small fraction of rubisco’s natural diversity has been kinetically characterized. Here, we present a large-scale kinetic survey covering the entire spectrum of rubisco’s diversity found in nature. Focusing on genetic clusters with above-average rates, we show that rubisco’s catalytic rate does not exceed ≈30 reactions per second at 30°C. Supported by a machine-learning predictive model, we extend this finding to all sequenced natural variants. Our study provides the most comprehensive kinetic dataset for a single enzyme to date.

## Introduction

Rubisco-based biological carbon fixation has shaped Earth’s atmosphere and biosphere (1, 2). It emerged about 3.5 billion years ago, during the Archaean, when carbon dioxide atmospheric concentration was >100,000 ppm and oxygen concentration was <1 ppm. One billion years later, oxygenic photosynthesis appeared in cyanobacteria, triggering the inversion of these two gasses’ atmospheric levels. Over geological time, atmospheric carbon dioxide and oxygen levels gradually changed to reach their pre-industrial levels of ≈280 and ≈200,000 ppm respectively. Catalyzing the main carboxylation reaction, rubisco is a central player in this history. Rubisco probably evolved in archaea, from closely-related isomerases called rubisco-like proteins (RLPs). Unlike RLPs, rubisco is able to catalyze the fixation of atmospheric CO_2_ into ribulose-1,5-bisphosphate (RuBP) to form two molecules of 3-phosphoglycerate (3PGA), which are subsequently used to build biomass. Rubisco evolved and diverged into four distinct forms across the different biological kingdoms: I, II, III, and II/III (thus named for its position between the previously discovered forms II and III). Form II, II/III, and III rubiscos consist of a homomeric core composed of two ≈50 kDa large subunits (L_2_) often forming higher-order oligomers. They are found in bacteria (form II, II/III & III), archaea (form II/III & III), and dinoflagellate algae (form II). Form I rubiscos additionally encompass a small subunit, interacting with the large subunit in a 1:1 ratio, and assembling into a heterohexadecameric structure (L_8_S_8_). They are found in bacteria but also in most photosynthetic eukaryotes like algae and plants. More recently, a clade sister of form I rubisco that evolved without small subunits has been identified and denoted form I’ (3).

Despite being a central enzyme shaping Earth’s biosphere and atmosphere for billions of years, rubisco is commonly seen as slow (4, 5), limiting the rate of carbon fixation (6) in some circumstances. With a median turnover number of ≈3 s^−1^ (interspecies median k_cat_) rubisco is much slower than the vast majority of central metabolic enzymes (5). Moreover, in the presence of oxygen, rubisco catalyzes a competitive oxygenation reaction that reduces the efficiency of its carboxylase activity even further. The scientific community has long tried to improve rubisco kinetic parameters (e.g. increasing the carboxylation rate and/or the affinity to CO_2_ relative to O_2_) in order to enhance carbon fixation *in vivo*. Yet these directed evolution efforts have met with only limited success (7–11), dampening hopes of making a better rubisco and raising an hypothesis that rubisco may actually have reached ‘Pareto optimality’ in which neither kinetic parameter can be further improved without negatively affecting the others (12–15).

However, out of the tens of thousands of rubiscos that have been sequenced so far, less than 200 are kinetically characterized in the scientific literature. Therefore, the above observations and interpretations were made based on a small and non representative fraction of rubiscos.

Here we present a wide systematic survey of carboxylation rates aiming to test the existence (or absence) of fast rubiscos in nature using a much larger and more representative dataset. We set up an experimental pipeline for the high-throughput expression and carboxylation measurements of hundreds of representative variants systematically spanning the unexplored rubisco diversity in the biosphere. We found that most rubisco variants across the tree of life are slow (k_cat,C_ < 10 s^-1^ at 30°C), and only a few of them show carboxylation rates in the range of 15-30 s^-1^. A deeper kinetic exploration of the somewhat faster rubisco subclades confirms the long standing hypothesis of a kinetically constrained enzyme now validated across the wide diversity in the tree of life. We show that the presence of relatively fast rubiscos mostly coincides either with a microaerobic environment, or with the presence of a CO_2_ concentrating mechanism. This observation argues in favor of the hypothesis of a trade-off between velocity and substrate (CO_2_) affinity in rubisco. Finally, we used machine learning approaches to study and predict rubisco carboxylation rates from this dataset, predicting rates for the ≈68,000 rubisco variants sequenced to date. Such work can help future engineering efforts aimed at fine-tuning rubisco’s catalytic efficiency for enhanced carbon fixation.

## Results

### Rubisco’s diversity across the tree of life

A thorough exploration of genomic and metagenomic data identified ≈68,000 unique rubisco sequences (see Materials and Methods). We performed annotation of a set of rubisco sequences clustered at 90% identity, revealing the four main forms of rubiscos (i. e. I, II, II/III, and III), in addition to the extra form I’ rubisco clade (3). These were visualized using uniform manifold approximation and projection (UMAP), a dimension reduction technique that projects rubisco’s sequence space into two dimensions while preserving local relationships (Fig. 1A).

**Fig. 1.**
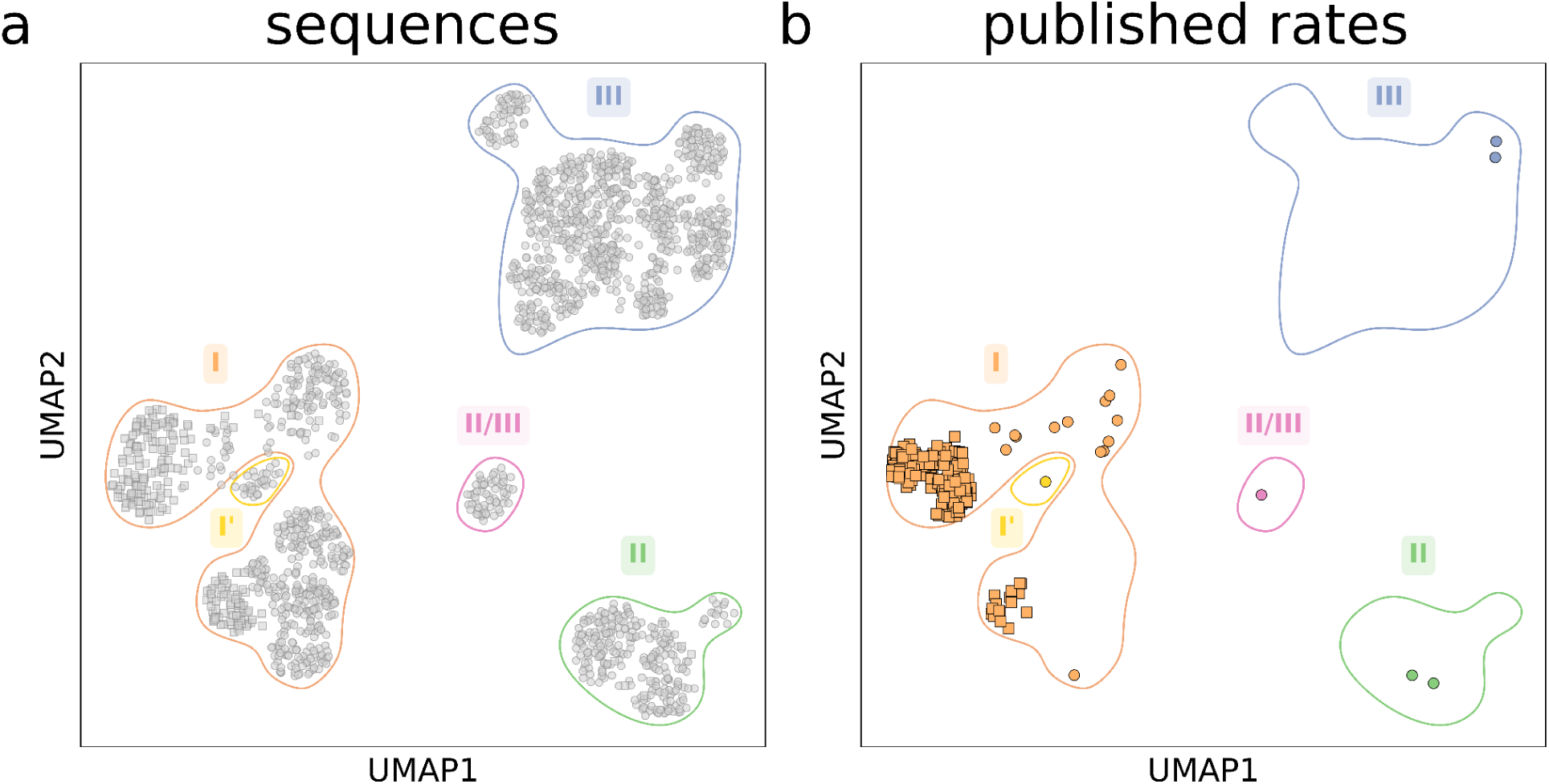
Rubisco’s natural diversity is mostly uncharacterized in terms of carboxylation rates. (A) Uniform manifold approximation and projection (UMAP) plot representing eukaryotic (squares) and prokaryotic (circles) rubisco natural diversity using a 90% sequence identity clustering. In (B), variants for which a carboxylation rate has been published in the literature before our work are highlighted, showing the sparse and highly biased coverage of the natural diversity.

With 61,000 rubisco homologs, eukaryotic form I rubiscos represent the most sequence-rich yet almost least-diverse rubisco clade: when plotted on a sequence distance basis, the sequence space of this over-represented group of rubiscos is comparatively narrow. Form II, II/III and III rubiscos are less numerous (850, 150 and 1600 sequences respectively) but present higher relative genetic diversity (Fig. 1A). This result is probably caused by a sampling bias, with over-representation of eukaryotic (plants and algae) rubiscos and poor sampling of less-accessible prokaryotic rubiscos. In order to quantify how well the subset of characterized rubiscos covers the full diversity, we first define a *representative set of sequences* based on a 90% identity clustering (in order to eliminate the sampling bias). These representatives are the sequences shown in the UMAP in Fig. 1A. Then, we define a *diversity coverage* metric for this study as the fraction of these representative sequences that are covered by characterized variants at a given identity threshold, X (i. e., that share at least X% sequence identity with any characterized variants).

We compiled all published carboxylation rates from prior studies that used active site quantification, a method that allows for precise measurements of turnover rates while considering the enzyme’s activation state, and plotted them together (Fig. 1B and Supplementary Data 1). With only 20 published rates, prokaryotic rubiscos have so far remained poorly explored: in terms of diversity coverage as defined earlier, rates available in the literature cover only 14% of them at an 80% identity threshold (Fig. S1A). The sequence space of prokaryotic rubiscos greatly exceeds their eukaryotic counterparts, and the diversity of these prokaryotes’ biotopes could have promoted the evolution of a more diverse range of carboxylation kinetics. Prokaryotic rubiscos thus represent a diverse, yet almost unexplored, domain of rubisco variety in the biosphere.

In contrast, most measured carboxylation rates available in the literature are from eukaryotic form I rubiscos. We observe that the rates available in the literature cover 87% of eukaryotic form I rubiscos at 80% identity (Fig. S1B). Additionally, the measured carboxylation rates have a range between 4-6 s⁻¹ (0.25-0.75 quantiles, when corrected to 30°C by assuming a Q_10_ value of 2.2 (16)). This low variation in carboxylation rates probably reflects the relatively low heterogeneity of the evolutionary habitats of these eukaryotic enzymes (mostly mesophilic, aerobic biotopes). Our efforts here therefore concentrated on the exploration of the uncharted rubisco diversity, specifically focusing on all prokaryotic rubiscos and a few non-type I eukaryotic rubiscos.

### High-throughput screening of prokaryotic rubisco carboxylation rates

As rubisco has long been posited to be a slow enzyme, our approach aimed at performing a high-throughput carboxylation survey in search of higher CO_2_ fixation kinetics. For each form, we chose sets of rubisco homologs whose number and identity threshold were adjusted to cover the sequence diversity of that rubisco form. We selected sets of 144, 18, 132, 24, and 105 form I, I’, II, II/III and III rubiscos respectively (see Materials and Methods), significantly improving the coverage of rubisco diversity in a systematic and comprehensive manner (Fig. S2). The representative sequences were codon-optimized for expression in *E. coli*, synthesized, and cloned into a pET vector system (see Materials and Methods, (17, 18)).

Expression and sample preparation were adapted to each rubisco form. For form II and II/III rubiscos, the simple homodimeric structure and, in the case of form II rubiscos, the fact that they mostly originate from *Pseudomonadota* (also known as Proteobacteria), like *E. coli*, facilitated relatively easy expression, without the need for any additional molecular chaperone. Moreover, rubiscos were expressed with an N-terminal His-tag followed by a SUMO protease recognition site, allowing for scarless cleavage-based elution, and resulting in purified native rubiscos (17). Over 70% of rubiscos were expressed and soluble with this protocol.

To avoid loss of enzyme subunits during the purification process, form I, I’ and form III rubiscos were not purified: kinetic measurements were performed directly from soluble protein extract. Additionally, individualized treatments were adapted to these rubisco forms in order to maximize the expression in *E. coli*. Form I rubiscos harbor a heterohexadecameric structure and originate from a more diverse set of bacteria phyla (including *Proteobacteria*, *Firmicutes*, *Chloroflexi*, *Actinobacteria*, *Calditrichaeota*, *Verrucomicrobia*, and *Cyanobacteria*). Coexpression with a defined set of chaperones (GroEL-GroES in addition to RbcX and/or Raf1 for β-cyanobacteria) allowed soluble expression of almost 80% (112 out of 144) of form I rubiscos in *E. coli* (18). Form I’ rubiscos were coexpressed with GroEL-GroES with a soluble expression rate close to 70% (12 out of 18).

Form III rubiscos were the most challenging variants to characterize. They were also coexpressed with the chaperone GroEL-GroES but showed a soluble expression rate of about 30%. Preliminary attempts to increase solubility by coexpressing archaeal chaperones were unsuccessful. This notably low expression, and the low measured carboxylation rates of the few soluble variants show the limit of the current pipeline for the study of such phylogenetically distant enzymes. The temperature, medium composition, and/or cellular machinery might be some of the factors present in *E. coli* that likely do not adequately support effective expression of these enzymes.

Yet, overall, these methods enabled us to measure the carboxylation rates of 221 representative rubisco variants using a coupled assay (Fig. 2; see Fig. S3 for a classic phylogenetic tree representation of all measured and published rates). Because coupled assays are known to underestimate rates compared to direct assays used in the literature, we applied a multiplicative correction factor based on a log–log comparison of rates from 11 rubisco variants measured in both our pipeline and the literature (See Materials and Methods, Fig. S4, and Table S1).

**Fig. 2.**
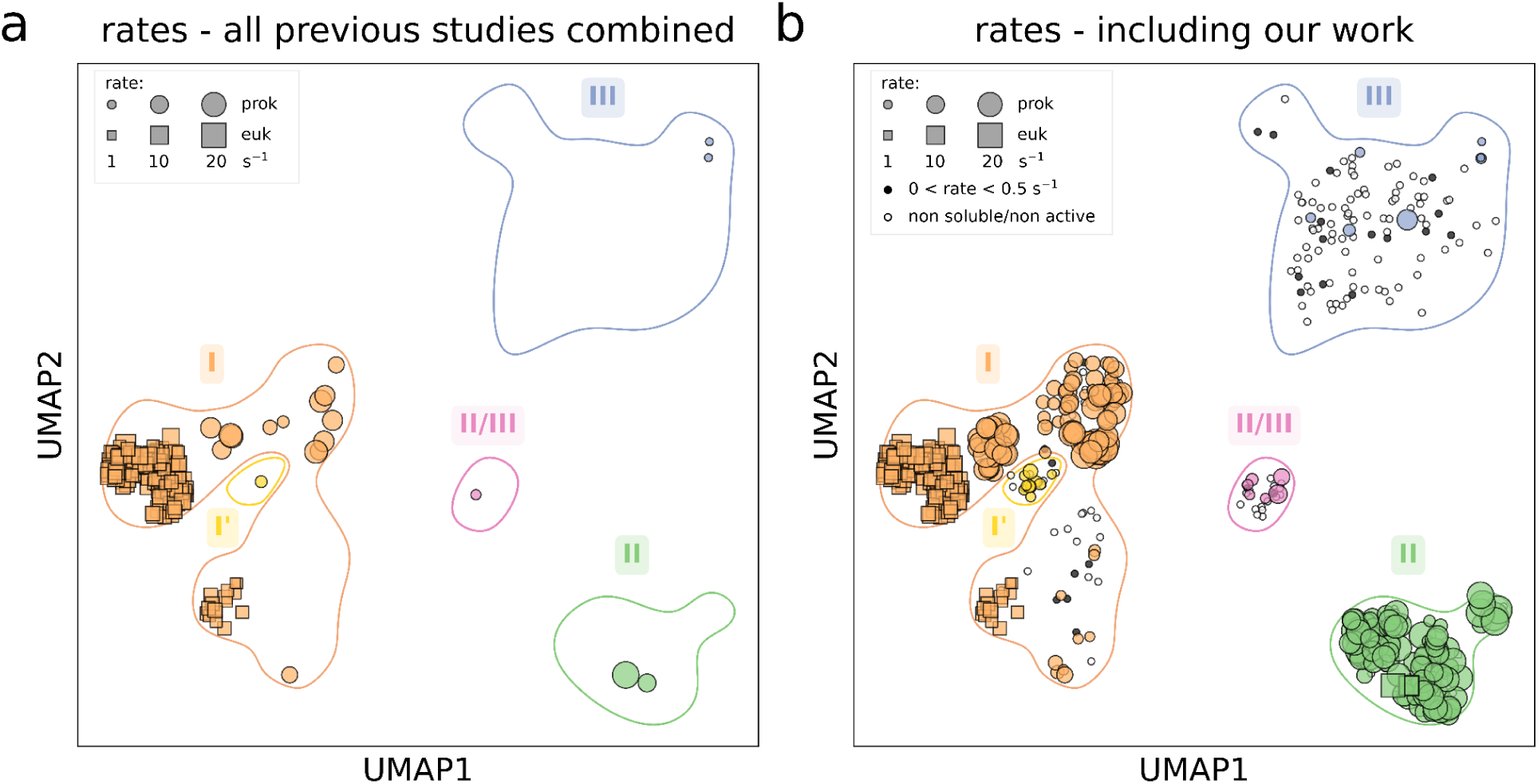
Systematic exploration of rubisco carboxylation rate covers a much larger fraction of the natural genetic diversity. UMAP plot representing rubisco natural diversity. Rubisco variants with a carboxylation rate reported in the literature (A) and in our work (B) are highlighted with size proportional to measured carboxylation rates. Rubisco variants that were either insoluble or inactive are represented as a white circle. Variants with carboxylation rates below 0.5 s^-1^ are represented as a black circle. All rates presented correspond to measurements or corrections made at 30°C.

We find that carboxylation rates of rubiscos from all forms are comparable and relatively low (Fig. 2B and S5 and Supplementary Data 2-3). Combining the two datasets—from our measurements (mostly prokaryotic variants), and previously published works (mostly eukaryotic rubiscos)—shows a complementary pattern in the distribution of measured rates (Fig. S5A). The median carboxylation rate of rubisco is 7 s^-1^, an order of magnitude lower than other central metabolic enzymes which typically harbor turnover numbers of ≈20-80 s^-1^ (5). This finding tends to confirm that, somewhat paradoxically given its central importance in the global food chain, rubisco is probably one of the slowest central metabolic enzymes in the biosphere.

### Deep exploration of faster than average clades does not reveal further enhancements in the carboxylation rate

Despite the relative low variation in measured carboxylation rates, we were able to characterize a few variants, among form I and II rubiscos, that harbored rates above average. Namely, within form I rubiscos, the highest carboxylation rates were observed in carboxysome-associated rubiscos, as seen in our previous work. An in-depth exploration of these rubiscos confirmed that carboxysome-association is the strongest biological indicator for fast carboxylating rubiscos among form I enzymes (18). Among form II rubiscos, the 10 fastest variants in the high-throughput screen (previous section) mostly belong to two clades: a) a clade composed of the *Piscirickettsiaceae*, *Gallionellaceae*, and *Mariprofundaceae* families and b) a clade composed of members of the *Paracoccaceae* family (17). We decided to explore more deeply these two clades in order to continue to search for enhanced carboxylation rates and address the question of rubisco rate limits at a finer resolution. We therefore set out to express and test all rubisco variants that belonged to these two genetic clusters (Fig. 3). Among the 44 rubiscos tested in this additional screening, 38 were soluble and active *in vitro*. We found that they all showed rates similar to their representative variants (Fig. 3). These findings suggest that we might be reaching a plateau in discovering higher carboxylation rates.

**Fig. 3.**
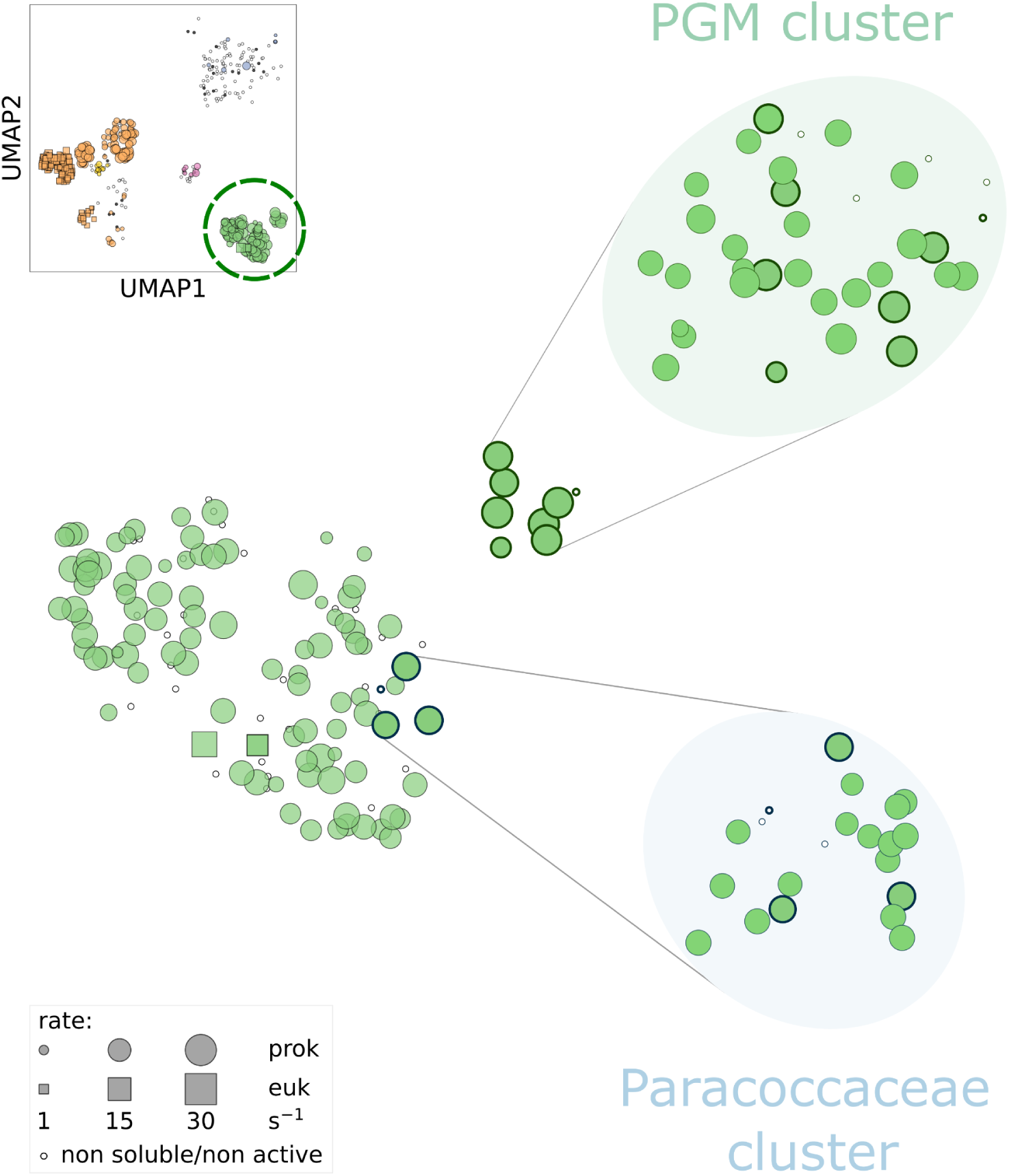
Deeper coverage of faster-than-average form II rubisco clusters does not show a significantly higher rate. UMAP plot representing the expansion of clades of fast carboxylating form II rubiscos (“Paracoccaceae” and “Piscirickettsiaceae, Gallionellaceae, and Mariprofundaceae” [PGM] genera clades). Original sequences from these expanded clades are represented with a thicker outer lining in both unzoomed and zoomed panels for clarity. Newly characterized rubiscos do not exceed 30 reactions per second.

Furthermore, analyzing sequence distances against carboxylation rate ratios among all tested rubisco variants in this work indicates limited rate variation (median rate ratio below 2) between sequences sharing 50% or greater identity (Fig. S6A). Moreover, when examining each specific rubisco form separately, a common trend is observed: closer sequences tend to have more similar carboxylation rates (Fig. S6B). Overall, our results suggest a relatively smooth rate landscape connecting functional rubiscos, meaning that among them, small sequence differences are unlikely to result in significant rate changes. The coverage achieved in our work (Fig. S2A) is likely comprehensive enough to support the conclusion that rubisco is generally slow across the tree of life.

### Reconstructed ancestors of faster-than-average form II rubisco clades are themselves among the fastest

In addition to an exploration of the extant natural sequence space level, we explored the two clusters of relatively fast carboxylating rubiscos by studying their ancestors. Enzyme ancestors have been noted to tend to harbor high thermostability and high catalytic activity compared to their contemporary descendents (19). We therefore reconstructed the ancestor of these two fast carboxylating rubiscos clusters. Both ancestors were soluble and active *in vitro*, and showed rates higher than the average rate of their contemporary descendents (Fig. 4). For the clade composed of the *Piscirickettsiaceae*, *Gallionellaceae*, and *Mariprofundaceae* families, the reconstructed ancestor even showed the highest rate (33.0 s^-1^), which makes it the fastest rate ever measured using our pipeline.

**Fig. 4.**
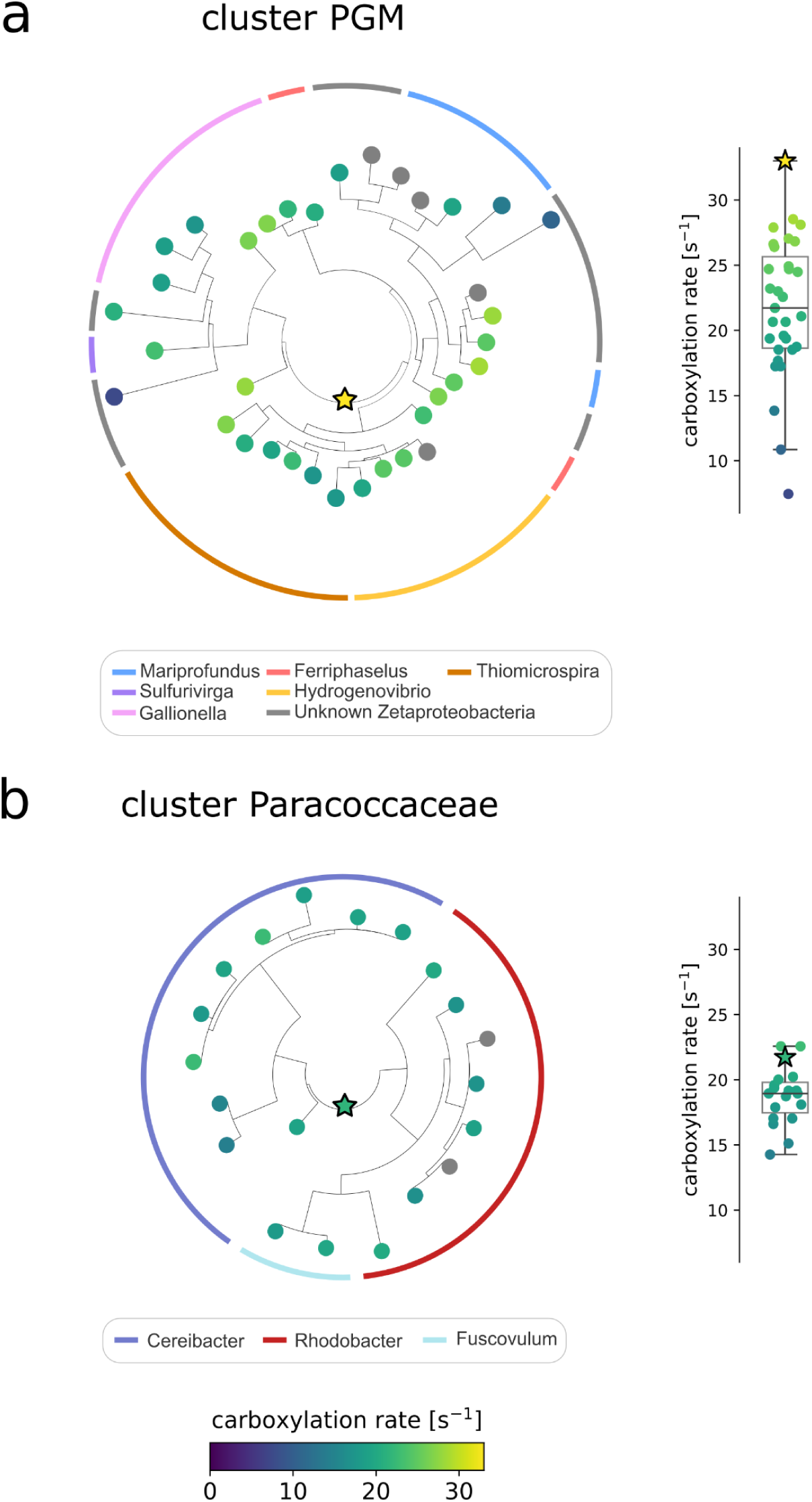
High carboxylation rates in reconstructed ancestors of faster-than-average form II rubisco clusters. Phylogenetic tree reconstructions of the two clades of faster-than-average carboxylating form II rubiscos: (A) “Piscirickettsiaceae, Gallionellaceae, and Mariprofundaceae” [PGM], and (B) Paracoccaceae. Contemporary rubiscos appear as circles and the reconstructed ancestors as stars. The carboxylation rates are represented by colors (between 0 and 33 turnovers per second). The same rates are shown also in a box plot on the right-hand side. The two external colored rings indicate the bacterial genus from which each rubisco originates.

These higher rates, confirming the expectations of high catalytic activity of reconstructed enzyme ancestors, are still relatively low when compared to other central metabolic enzymes, and reinforce the notion of a kinetically constrained enzyme.

### Machine learning approaches identify residue associated with rubisco carboxylation rate and predict uncharacterized rubiscos rates

This dataset can be used for machine-learning approaches aiming to link rubisco sequences and their associated rates, especially as similar sequences generally exhibit similar rates (Fig. S6). As a first approach, focusing on form II and II/III rubisco sequences, which show relatively high sequence conservation, we performed random forest regression to predict rubisco’s carboxylation rate association to the amino acids at each position (Fig. 5, see Materials and Methods). The derived Shapley additive explanations (SHAP) values (20, 21) quantify the association of each tested position with the predicted rate (Fig. 5A). Position 327 (position number based on *Rhodospirillum rubrum* rubisco) has a SHAP value above 1.2, representing an association between the amino acid at this position and the carboxylation rate. The presence of a tyrosine at this position is associated with a significantly higher carboxylation rate (Fig. S7 and 5B). This position is located in the mobile loop 6, which transitions between the open and closed state during each catalytic cycle (22). It is two residues upstream of K329, a conserved residue at the apex of this loop positioning the substrate CO_2_ for catalysis (Fig. 5C) (23, 24). The equivalent of F/Y327 in form I rubiscos (V332) comprises the loop 6 hinge (25) and its sidechain is anchored in a hydrophobic pocket. The hydroxyl group of tyrosine may increase the flexibility of the loop 6 hinge, and thus accelerate carboxylation velocity (permit more rapid cycling between open and closed states) when placed in the correct context. However, the possible epistatic effect of other amino-acids in the sequence makes it unlikely to be a mutational hotspot for improving rubisco carboxylation rate. Introducing a tyrosine at this position had a neutral or slightly positive effect in some variants, but led to a rate decrease in *Candidatus Peregrinibacteria* rubisco, probably due to other epistatic effects (Fig. S8).

**Fig. 5.**
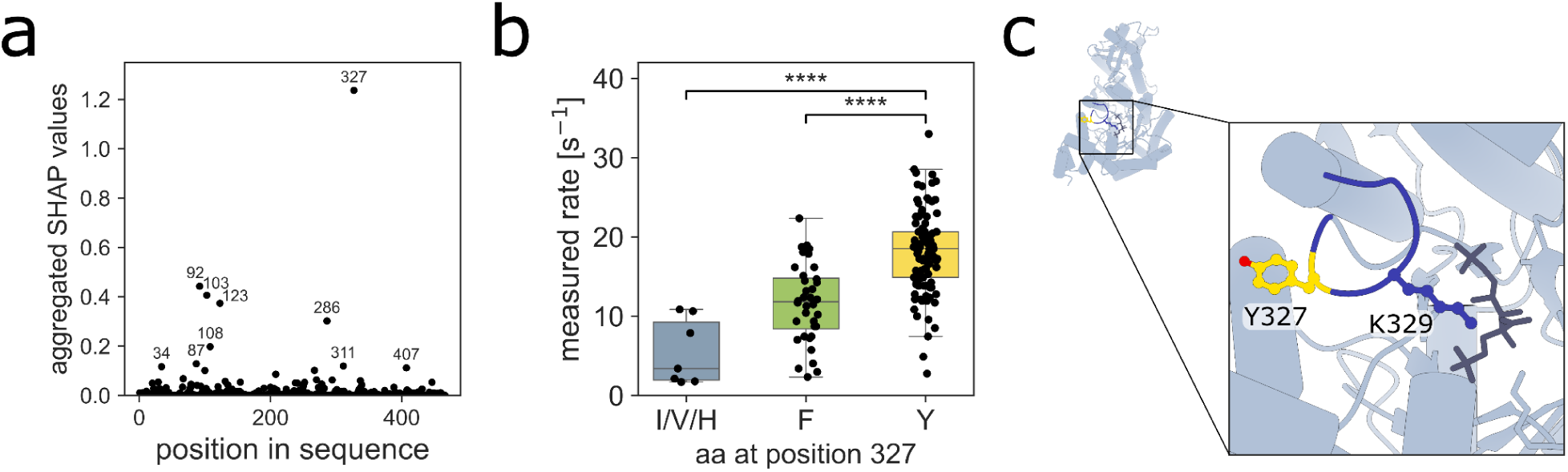
Random forest regression on form II and II/III rubisco variants shows residue 327 is associated with rubisco carboxylation rate. (A) Residue importance was determined using absolute SHAP (Shapley additive explanations) values from a random forest regressor model. The model assessed the rubisco carboxylation rate based on each amino acid. (B) Rubisco carboxylation rates as a function of the amino acid at position 327 (I/V/H: isoleucine/valine/histidine; F: phenylalanine; Y: tyrosine). (C) 3D structure of the CABP-bound rubisco from Gallionella highlighting Tyr327 (yellow) and Lys329 on loop 6 (dark blue). Kruskal-Wallis followed by Dunn multiple comparison tests were applied. ****p < 0.0001.

Another way to leverage the dataset of over 250 carboxylation rates from our work, along with the 190 rates from previous studies in the literature, is to use it as a reference for predicting the carboxylation rate of any rubisco from its sequence. To do so, we trained several simple machine learning models including a nearest neighbor model, unweighted and weighted mean models, and a support vector regression (SVR) model (26) (see Materials and Methods). We assessed the prediction quality with a leave-one-out cross-validation, using subsets sharing increasing identity with the sequence to predict (Fig. S9). In all models, the root mean square error (r.m.s.e.) of carboxylation rate predictions mostly decreased with increasing sequence identity thresholds, i.e., using characterized variants subsets that share increasing identity with the one to predict. The SVR model shows reliable performance, with significant improvement compared to taking the average rate of each subset, especially for identity thresholds above 80%. We therefore used it to predict the carboxylation rate of all ≈68,000 sequenced rubisco variants found in nature to date (Supplementary Data 4). We predicted the rates of more than 90% of the rubisco variants with less than a 2-fold change estimated error (defined as one standard deviation), achieving predictive performance comparable to other large-scale machine learning efforts linking protein sequences to catalytic rates (27–29). The model’s predictive accuracy was lower only for certain groups, such as the less-characterized form III rubiscos for which we only screened at an identity threshold of 55%, resulting in a ≈3 fold change r.m.s.e. All predicted rates range up to 26 s^-1^. We compared predicted rates across different rubisco forms (Fig. 6 and Table S2). To ensure precision, partial sequences (54,000 variants, mostly eukaryotic) were excluded from this analysis. With a median rate of 15 s^-1^, form II rubiscos are the fastest, ahead of any other forms, which have median rates ranging from 1.9 to 4.4 s^-1^. As a curiosity, a leave one out cross validation of the fast rubisco ancestor from the *Piscirickettsiaceae*, *Gallionellaceae*, and *Mariprofundaceae* families would indeed have predicted a rate of 24.6 s^-1^ placing it in the top 0.02% (rank 14 out of 67,706) when compared to predicted rates of the 67,706 rubisco unannotated variants (Fig. S10). Yet, in general, in silico results should be treated especially carefully for resurrected ancestries, as reconstructed or consensus variants tend to sit at the center of the family alignment and inherit an excess of stabilizing residues, leading most machine-learning methods, SVR included, to over-score them (30). Beyond providing the predicted rates of all sequenced rubisco variants in nature, this model establishes a critical benchmark for any future machine learning approach aiming to predict rubisco rates.

**Fig. 6.**
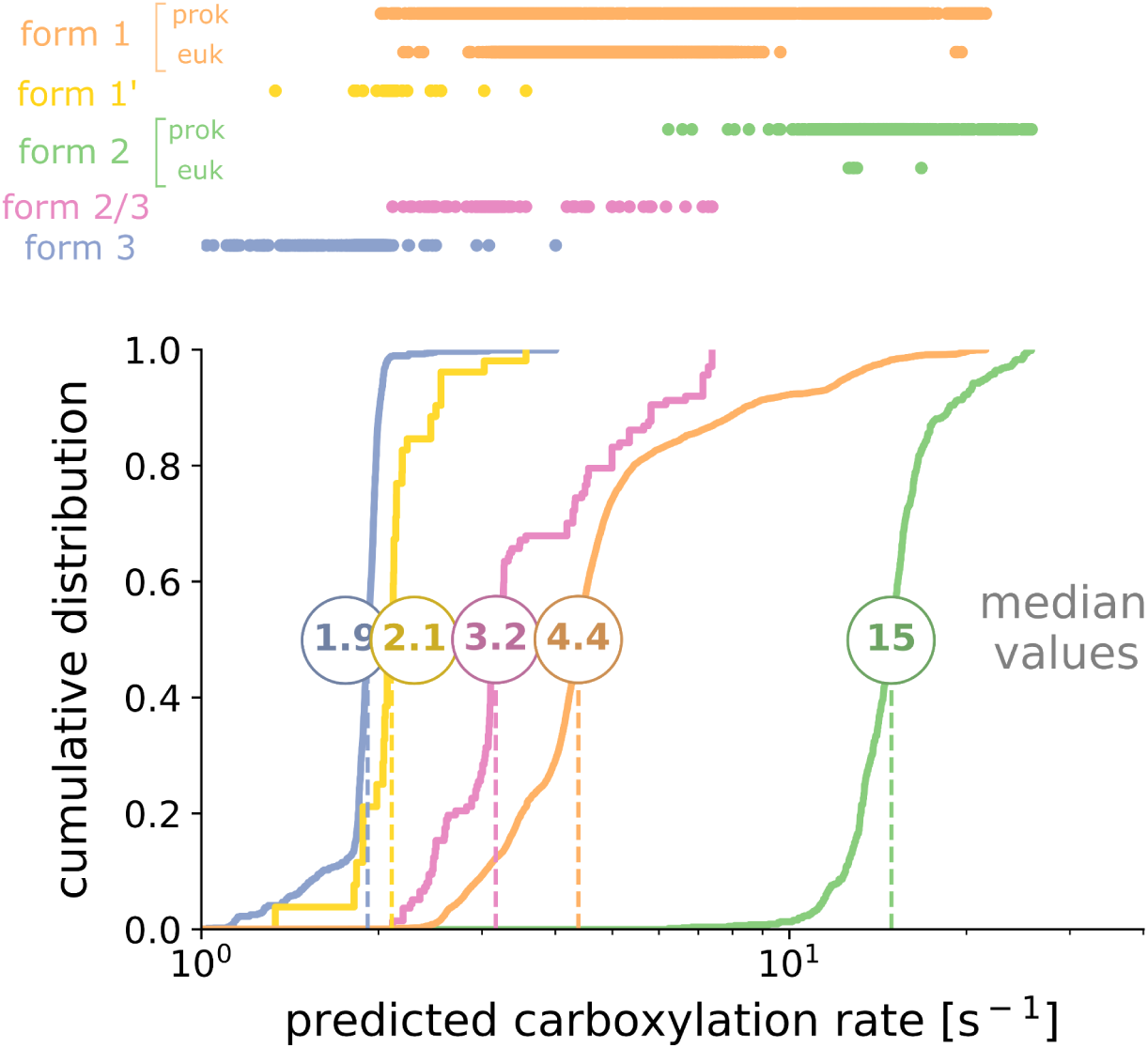
Prediction of carboxylation rates for rubisco variants found in nature using a support vector regression (SVR) model. Distribution of predicted rates is shown by rubisco form. Only the complete sequences were used for this plot.

## Discussion

Rubisco is an enzyme that has drawn much attention since its first discovery as “Fraction I” in 1947 (31). As it appeared early in the history of life and plays a central role in autotrophy, many consider it a marker of evolution across different ecological niches and the changing atmospheric composition. Moreover, its notoriety for being slow and a limiting factor for plant growth makes it a major target for improvement from plant scientists to synthetic biologists.

Found in the three domains of life—bacteria, archaea, and eukarya—it has evolved a wide sequence diversity that has so far been mostly unexplored, especially among prokaryotes. This paper reports a systematic investigation of this diversity by studying a central kinetic parameter, the carboxylation rate, and answering the question of whether much faster rubiscos exist.

Through the screening of ≈500 representative rubisco variants spanning the uncharted rubisco diversity, we bring evidence that rubisco is slow across its wide natural genetic diversity. Paradoxically with its evolutionary success as the primary catalyst for carbon fixation on Earth, this relative slowness likely reflects a chemical compromise: catalyzing the complex multistep reaction of CO_2_ fixation onto RuBP in an oxygenic atmosphere (for aerobic ones), and matching various metabolic demands (which can depend on the trophic mode, the availability of other limiting nutrients, etc.) (32). Therefore, rubisco has probably reached ‘Pareto optimality’ and shows a relatively average rate when compared to other enzymes. However, when compared to other central carbohydrate energy metabolism enzymes (from glycolysis/gluconeogenesis, the citrate cycle, pyruvate metabolism, etc.), with a median turnover of ∼79 s⁻¹ (5), and even to other carboxylases (33), rubisco stands as a relatively slow catalyst.

Form III rubiscos were the most challenging enzymes to study, likely due to the suboptimal conditions in our system. Furthermore, even the ones that were soluble had a very low carboxylation rate (median rate of ≈2 s^-1^) or showed no activity at all in our assay. Likewise, form II/III rubiscos have poor carboxylation kinetics (median rate of ≈3 s^-1^). This is possibly due to their metabolic context: they typically serve in nucleoside salvage (34, 35) or in carbon metabolism pathways distinct from the classical Calvin-Benson-Bassham (CBB) cycle (36–38). The evolutionary pressure exerted on these rubiscos may have been aimed towards objectives other than higher carboxylation rates. Finally, it is noteworthy that some of the rubiscos come from thermophilic organisms. These rubiscos could therefore display faster rates if tested at very high temperature, like the form III rubsico from *Archaeoglobus fulgidus* which has a k_cat,C_ of 23 s^−1^ at 83°C (39).

Other notably slow rubiscos were form I’ rubiscos (median rate of 2.1 s^-1^), or some subgroup of form I rubiscos, like the recently discovered “form I Anaero” (median rate of 3.7 s^-1^). Both of these rubisco groups are thought to have evolved in anaerobic conditions, before the appearance of cyanobacteria (3, 40). Interestingly, this could be seen as a contradiction with the hypothesis of a trade-off between rubisco carboxylation rate and CO_2_ affinity as the absence of oxygen could release the pressure towards high CO_2_ affinity. However such catalytic trade-off may stand only when there is an evolutionary pressure on rubisco/when CO_2_ fixation is a limiting factor. This is likely to depend on the organism and/or situation. These slower carboxylating rubiscos may thus be adapted to lower metabolic needs of anaerobic bacteria (in comparison to oxygenic phototrophs for instance). Also, it is currently unclear whether “form I Anaero” rubiscos are expressed by autotrophic or heterotrophic bacteria. The latter could be possible as the CBB cycle has for instance been reported to serve as a secondary electron sink in some heterotrophic bacteria (41). Rubisco enzymes involved in such functions may also experience less pressure towards the evolution of faster carboxylation rates, as compared to autotrophy-oriented rubiscos.

We report the presence of fast carboxylating rubiscos in the two remaining rubisco forms. Among form I, the fastest rubiscos were found to be associated with a carboxysome (18). Among form II, the fastest variants were mostly found within two monophyletic groups, which share the feature of mostly originating from microaerophilic bacteria (*Mariprofundus*, *Ferriphaselus*, *Sulfurivirga*, *Gallionella*, *Hydrogenovibrio* sp. SC-1), or bacteria with an at least facultative anaerobic lifestyle (*Rhodobacter*, *Cereibacter*, *Fuscovulum*). As discussed in de Pins *et al.* 2024, these observations could support the hypothesis of a trade-off between rubisco carboxylation rate and CO_2_ affinity: rubiscos evolving in the context of an elevated CO_2_/O_2_ ratio may have greater ability to evolve towards high carboxylation rates. Interestingly, the few relatively fast form II rubiscos expressing bacteria that were reported to be strictly aerobic (*Thiomicrorhabdus aquaedulcis*, *indica*, and sp. Kp2, *Hydrogenovibrio marinus*, *kuenenii*, and sp. Milos-T1, *Thiomicrospira* sp. XS5) all express an additional rubisco. As we suggested in Davidi *et al.* 2020, these form-II variants are likely active under low oxygen conditions (42–44).

In addition, we observed that rubiscos sharing similar sequences tend to have closer rates (Fig. S6A), and that this trend seems to hold within each rubisco form (Fig. S6B). Considering the strength of this phylogenetic signal is important to better evaluate the real constraints imposed by catalytic trade-offs such as the affinity/velocity trade-off. An important work from Bouvier and colleagues began to address this question in rubiscos from phototrophs (45); our expanded dataset could therefore serve as a basis for measurements of additional kinetic parameters (e.g., CO₂ affinity and specificity) and further enrich models that aim to accurately determine the trade-offs constraining rubisco kinetics (45–47).

Lastly, we note that these fast rubisco monophyletic clades do not strictly follow the phylogeny of their bacteria, i. e. some rubiscos sharing high sequence identity are expressed by relatively distant bacteria phylogenetically (*Piscirickettsiaceae*, *Gallionellaceae*, and *Mariprofundaceae* families represented in the fastest form II rubisco monophyletic clade for instance). This could suggest a convergent evolution of these genes, or horizontal gene transfers between distant species: bacteria adapted to analogous selective pressures (like a high CO_2_/O_2_ ratio) to develop, transfer, and/or maintain highly similar rubisco genes.

To further explore carboxylation rate limits, at a finer resolution, we systematically screened all variants belonging to two clades of relatively fast form II rubiscos. In this second round of screening, we did not measure any rate exceeding 30 s^-1^. This strongly advocates for a highly constrained variation of this kinetic parameter. The fastest rubisco measured through our pipeline appears to be the reconstructed ancestor of one of the relatively fast form II rubisco clades. Future research could explore more extensively, not across the “natural sequence space” axis, but on a “temporal” axis, across all rubisco ancestors, as some studies have begun doing (8, 40, 48–51).

Note that in the literature, the direct radiometric rubisco assay yields highly variable carboxylation rate values for the same variants, as analyzed in detail by Iñiguez and colleagues (49). We aimed here to reconcile spectrophotometric and radiometric assays by applying a correction factor based on the comparison of values obtained for the same enzymes. Further work may be needed to accomplish this with higher precision, but we believe the strength of our current pipeline lies in its systematic approach, in particular the consistent inclusion of a control rubisco in every assay (here, the form II variant from *R. rubrum*). We propose that the systematic inclusion of a reference control in future kinetic studies would help advance this objective.

This extensive dataset of rubisco sequence representatives and their associated carboxylation rates can further be used to link sequence motifs to catalytic function. A random forest regression on form II and II/III sequences identified one such position associated with rubisco carboxylation rate. This position belongs to the mobile loop 6 of rubisco, a loop known to be involved in catalysis (24), and thought to stabilize the reaction intermediates. In tobacco rubisco, during the transition from the open to the closed conformation, the valine at this position undergoes changes in phi and psi backbone angles, while its sidechain remains in place (25). Making this sidechain more hydrophilic, with the hydroxyl group of a tyrosine for instance, could possibly influence this transition by enhancing loop mobility. Furthermore, work by Okano *et al* described that in the red alga *Galdieria partita*, this same valine is involved in a main chain oxygen hydrogen bond with a glutamine residue in helix α7, stabilizing the loop 6 in the closed state of the enzyme and thereby contributing to a higher CO_2_ affinity (52). This position could therefore be involved in the aforementioned affinity/velocity trade-off in rubisco. However, improving the carboxylation rate from single point mutations is notoriously difficult (53), as confirmed by our directed mutagenesis experiments which did not show any enhancement of the rate. Further work could identify whether additional mutations are necessary and could increase rubisco carboxylation rate.

We ultimately used this dataset as a reference to predict the carboxylation rate of all other uncharacterized rubiscos. Predictions match experimental data, with form II variants showing higher rates than other rubisco forms. As already mentioned, certain groups, such as form III rubiscos, underwent less stringent screening (identity threshold of 55%), resulting in a ≈3 fold change r.m.s.e in the predictions. However, as the slowest characterized rubiscos, these groups are unlikely to harbor unusually fast outliers or challenge the hypothesis of a globally slow rubisco across the tree of life. Interestingly, among eukaryotic form I rubiscos, three variants stand out from the rest of the group, with rates close to 20 s^-1^, more comparable to the rates of fast prokaryotic form I variants. These variants are found in photosynthetic amoeboids of the genus *Paulinella*, which result from the recent endosymbiosis of an α-cyanobacterium in these originally heterotrophic species (54). In contrast, most other photosynthetic eukaryotes trace their origin to a primary endosymbiosis involving a β-cyanobacterium ancestor (55). Alpha-cyanobacteria are known to express some of the fastest carboxylating rubiscos among form I enzymes (18), which likely explains the elevated catalytic rates observed in *Paulinella* variants. Yet, overall, our machine learning approach supports the hypothesis that rubisco is slow across its genetic diversity, with no predicted rates exceeding 26 s^-1^. Moreover, it enriches ongoing efforts to apply machine learning methods to predict enzymatic rates (27–29). In particular, Iqbal and colleagues employed Gaussian processes (GPs), a Bayesian machine learning method, to predict plant IB rubisco kinetics from its large subunit sequence (56). Our study offers a complement to this work, as we focus on a much broader sequence space, including all microbial rubiscos, leveraging our expanded dataset of experimentally measured rates. While their model shows strong predictive performance, it was restricted to a narrower group of rubiscos, which naturally facilitates lower prediction errors. Our aim here was to apply a basic machine learning algorithm, as a baseline benchmark for future studies aiming to predict rubisco carboxylation rates, and providing a reference point against which the performance of more advanced and promising algorithms (57) can be evaluated.

The work reported here systematically addresses the question of how much the previously uncharacterized genetic diversity of rubisco holds enhanced carboxylation rates. The experimental strategy we developed for this purpose led us to screen a large cohort of ≈500 representative rubisco variants covering much of the remaining diversity in the tree of life not explored previously. Leveraging machine learning models, we extrapolated these findings to encompass all currently known diversity of this enzyme. We demonstrate here that rubisco is relatively slow across this wide variation in the tree of life.

## Materials and Methods

### Rubisco kinetic data collection

Carboxylation rates reported in the literature were compiled, corrected to 30°C considering a Q_10_ value of 2.2 (16), and the median was taken (Supplementary Data 1). Only values coming from CABP-based active site quantification were considered, with the exception of the rates from the two form III rubiscos (58, 59). This exception was made due to limited kinetic data available for this rubisco form.

### Rubisco sequence collection and variants selection

Thorough exploration of genomic and metagenomic data led to the identification of ≈68,000 unique rubisco sequences. Essentially, rubisco large subunit sequences were collected from NCBI’s nr database downloaded in December 2020, in-house assemblies of samples from Tara Oceans expeditions (60) and various published assemblies and sequences (3, 61–64). Sequences shorter than 300 amino acids or longer than 700 amino acids were filtered out, leaving a set of ≈72,000 sequences. Sequences were then clustered at 80% identity using USEARCH algorithm (v8.1.1861_i86linux64, parameters-cluster_fast-id 0.8) (65). Cluster representatives were then aligned with MAFFT (v7.475, default parameters) (66) and columns with more than 95% gaps were removed using trimAl (v1.4.rev15,-gt 0.05) (67). Rubisco forms were identified through a phylogenetic tree constructed using FastTree (v2.1.10, default parameters) (68), and annotated based on existing data from NCBI, (69), and (3). ≈5,000 non-carboxylating rubisco-like proteins (RLPs) were identified and filtered out, resulting in a set of ≈68,000 unique rubisco homologs.

For rubisco variant selection, a sequential screening approach based on rubisco large subunit sequence clustering, at varying sequence identities, was adopted. For form I variants, the approach was already outlined in (18). For form I’ and III variants, representatives of clustering at 85 and 55% identity were respectively selected to cover these two rubisco forms with a number of variants (18 and 105 respectively) we could afford to synthesize in the span of this study. For form II and II/III variants, all 143 representatives of clusters at a 90% identity threshold were screened, along with 2 control variants as previously described in (17). These were supplemented by 11 other rubiscos arbitrarily selected for data completeness. After identification of two monophyletic groups of fast rubiscos (composed of 8 and 4 rubiscos respectively), we further expanded these groups to screen all (44) variants within them, based on the rubisco sequence dataset available to us at that time (November 2019). Ancestral rubisco variants were subsequently reconstructed for both groups (see below). In total, 180 new rubiscos were screened in this work, in addition to the 289 variants previously analyzed in our two earlier studies (17, 18).

### Gene synthesis

The selected rubisco genes were codon optimized for expression in *E. coli* (Twist Codon Optimization tool), synthesized, and cloned into an overexpression vector by Twist Bioscience.

For form I, I’, and III rubiscos, the vector used was pET-29b(+) (NdeI_XhoI insertion sites). For form II and II/III rubiscos, the vector used was a custom pET28-14His-bdSumo vector, as described in (17), allowing the fusion of a 14xHis-bdSumo tag to rubisco large subunit. Validation of gene synthesis and cloning was conducted through next-generation sequencing by Twist bioscience.

### Rubisco expression and preparation

Rubisco expression and sample preparation was performed as described in (18) for form I, I’, and III rubiscos, and as described in (17) for form II and II/III rubiscos. Essentially, plasmids were transformed into chemocompetent BL21(DE3) cells, priorly transformed with a chaperone-expressing pESL plasmid in some cases: the chaperone GroEL-GroES (70) for form I, I’, and III; or GroEL-GroES in tandem with the chaperone rubisco accumulation factor 1 (Raf1) for some insoluble form IB rubiscos (18). Transformed cells were grown at 37°C, 250 rpm in 8 ml of LB media supplemented with 50 μg/ml kanamycin and/or 30 μg/ml chloramphenicol (depending on the presence of the pESL plasmid for chaperone expression) in 24-deep-well plates. When cells reached an OD_600_ of 0.6, GroEL-GroES and/or Raf1 expression was induced by adding arabinose (0.2% final) and incubating at 23°C for 45 min. Rubisco expression was then induced by adding 0.2 mM IPTG (isopropyl β-d-thiogalactoside) and incubating at 23°C for 21 h. In the case of form II and II/III rubiscos, expression was simply induced when cells reached an OD600 of 0.8, by adding 0.2 mM IPTG and incubating at 16°C for 16 h. Cells were then harvested by centrifugation (15 min; 4,000 g; 4°C) and pellets were lysed with BugBuster® ready mix (Millipore) (70 µl for form I, I’, and III/ 500 µl for form II and II/III rubiscos) for 25 min at room temperature. Crude extracts were then centrifuged for 30 min at 4,000 g, 4°C to remove the insoluble fraction.

For form II and II/III rubiscos, a subsequent purification step of the enzymes was performed. The soluble fraction was transferred to 96-deep-well plates together with 100 µl of Nickel magnetic beads (PureProteome™; Millipore), following the manufacturer’s protocol for washing and binding. After 3 washes, 150 μl of cleavage buffer (20 mM EPPS pH 8.0; 50 mM NaCl; 20 mM MgCl_2_; 15 mM imidazole) containing bdSENP1 protease (8 μg/ml) was added to each well and incubated for 1 hour on a plate shaker to cleave the SUMO tag and elute rubisco. Purified proteins were then separated from the tag-bounded magnetic beads using a magnetic rack. Protein concentrations were measured using a Pierce™ BCA Protein Assay Kit (Thermo Fisher Scientific) according to the manufacturer’s multi-well plate protocol.

For quality control, all samples (0.2 μl of the crude extracts and 2 μl of the soluble fractions for form I, I’, and III/ 10 µl of the purified fraction for form II and II/III rubiscos) were run on an SDS–PAGE gel.

### Kinetic measurements

The carboxylation rates of each rubisco were measured in a high-throughput spectrometric coupled assay (as described in (17, 18)). Essentially, 3-phosphoglycerate production by rubisco was coupled to NADH oxidation, which decay can be measured spectrophotometrically, thus allowing inference of the specific activity of the sample. To minimize competitive oxygenation reaction, the assays were performed in a solution equilibrated at 4% CO_2_ (100 times atmospheric conditions) and 0.4% O_2_ (2/100 of atmospheric conditions) in a gas-controlled plate reader at 30°C. Rubisco’s active site concentration was measured in each sample by repeating the assay with increasing concentrations of 2-C-carboxyarabinitol 1,5-bisphosphate (CABP), a competitive irreversible rubisco inhibitor. Practically, for form II and II/III variants, as described in (17), the purified rubiscos were added to the reaction mix at a final concentration of 80 nM. For other rubisco forms, as described in (18), kinetic assays were performed directly from the soluble fraction of prepared lysates. Because this method does not allow for an estimation of rubisco concentration *a priori*, an initial assay was performed with the undiluted soluble fraction. When needed, i. e. when the rubisco was too concentrated to measure a decay of the activity with CABP, the assay was repeated with a diluted soluble fraction.

Rubisco carboxylation rates were eventually obtained by dividing the specific activity of each sample by its active site concentration. To account for variability across all assays, *R. rubrum* rubisco was consistently measured as a standard, and used to normalize all rates measured through our pipeline. Since the coupled assay used in our pipeline tends to underestimate the rates, the obtained rate values were finally corrected by scaling them with a multiplicative factor of 2.1 (95% confidence interval: 1.7-2.8), derived from a log–log comparison (slope fixed to 1) of rates from 11 rubisco variants measured in both our pipeline and the literature (Fig. S4). Note that measurements presented in this work were in part published in (17, 18), especially for form I, form II and II/III rubiscos. However, values presented in this work differ due to the normalization method employed. All individual measured rates before correction are provided in Supplementary Data 2, while the final processed rates used for analysis are in Supplementary Data 3.

### Rubisco sequence diversity plotting

To represent rubisco sequence diversity across all forms, the entire dataset of rubisco large subunit sequences was clustered at 90% identity. Rubisco sequences of variants that have been measured for their carboxylation rate in previously published work, or in our work, were also added to the dataset. All sequences were then aligned and a distance matrix was computed using Clustal Omega (71, 72). Multidimensional scaling (MDS) was performed to convert the distance matrix into a 6-dimensional vector space and a uniform manifold approximation and projection (UMAP) was subsequently applied to reduce the dimensions to 2. The specifics of the above procedure have been heuristically determined only for producing a desired visual effect, and have no bearing on the statistical analyses or predictions of turnover values in this work.

In cumulative plots estimating rubisco diversity coverage (Fig. S1 and S2), coverage is defined as the fraction of representative sequences, from a 90% sequence identity clustering, that are covered by at least one kinetically characterized rubisco variant at a given identity threshold, X (i. e., that share at least X% identity with any characterized variants). This approach quantifies the coverage of sequence diversity while minimizing biases from the over-representation, in number of sequences, of certain genetic groups, such as eukaryotic form I rubiscos, which may be due to sampling preferences.

### Ancestor reconstruction

The sequences from the two identified clades of fast carboxylating rubiscos were aligned, and phylogenetic trees were constructed using PhyML (73). Trees were midpoint rooted and used as inputs for ancestral sequence reconstruction using codeml from the PAML package (74). The JTT evolutionary model and marginal probabilities were used. For further testing, the sequences generated from the highest posterior probabilities at sites were selected.

### Random forest regressor model and feature importance analysis

A random forest regressor model was trained on the 148 characterized form II and II/III rubisco variants to predict rubisco carboxylation rate as a function of its sequence. These rubiscos’ sequences were aligned, one-hot encoded into a numerical format, and randomly split into training (75% of the data) and testing (25%) sets 100 times. For each iteration, an individual decision tree was trained with a maximum depth of 3, and the Shapley additive explanations (SHAP) values of each estimator (i. e. each aminoacid that can be found at each sequence position) were computed. The combined SHAP values at each position were eventually averaged and plotted together to compare the contribution of each position to the difference between the model’s prediction and the measured value.

### Machine learning models and rate prediction

We combined carboxylation rates from our work with those from previous literature, as training data for predicting the carboxylation rate of any rubisco from its sequence. We aligned the ≈68,000 rubisco’s large subunit sequences, dropping alignment positions that contained gaps in more than 95% of sequences. We computed sequence identity using Hamming distances between the aligned sequences. As features, we used a one-hot-encoding of the aligned sequences. We trained machine learning models using the 440 sequences with known rates. Predictions for each rubisco were made using only training data from the same form. Since carboxylation rates are approximately log-normally distributed per form (Fig. S5B) and it is standard practice in rate predictions (75–77), we log-transformed the carboxylation rates using the natural logarithm prior to training the models. We evaluated several models with a leave-one-out cross-validation using training sets of variants with increasing identity to the variant being predicted. These models included (i) nearest neighbor (NN), which predicts rates based on the most similar sequence in the training set; (ii) unweighted mean, which uses a simple average of the rates from this same set; (iii) weighted mean, which calculates an identity-weighted average using exponential weights based on sequence identity to the predicted variant; and (iv) support vector regression (SVR) with a radial basis function (RBF) kernel, which learns a nonlinear mapping between sequences from the set and their rate values. To assess the performance of each model we computed the root mean squared error (r.m.s.e) for each identity threshold using leave-one-out cross-validation (Fig. S9). Since SVR was the best performer in nearly all thresholds tested, we therefore then trained separate SVR models for each rubisco form using the full dataset of 440 characterized sequences, and used them to infer the rate of all ≈68,000 sequenced rubisco variants found in nature to date. For all validations and predictions, we used the scikit-learn package (78) with the default parameters for SVR (see “data and code availability” for details).

## Data and code availability

All data and code used for the analysis presented in this paper are open source and available in the following link: https://gitlab.com/milo-lab-public/rubisco_is_slow.

## Acknowledgements

We thank Yoav Peleg, Ilya Kublanov, Noam Prywes, and Avi Flamholz for their constructive feedback and insights throughout this work. We thank Zhijun Guo and Yi-Chin Candace Tsai for producing RuBP and CABP. This research was supported by the Mary and Tom Beck Canadian Center for Alternative Energy Research, Miel de Botton, the Schwartz Reisman Collaborative Science Program, and the Charles and Louise Gartner Professorial Chair. C. M. and A.-F. B. thank the European Research Council (ERC) for funding under the European Union’s Horizon 2020 research and innovation program (grant agreement no. 851173, to A.-F. B.).

