## Supplementary Information for "Rubisco is slow across the tree of life"

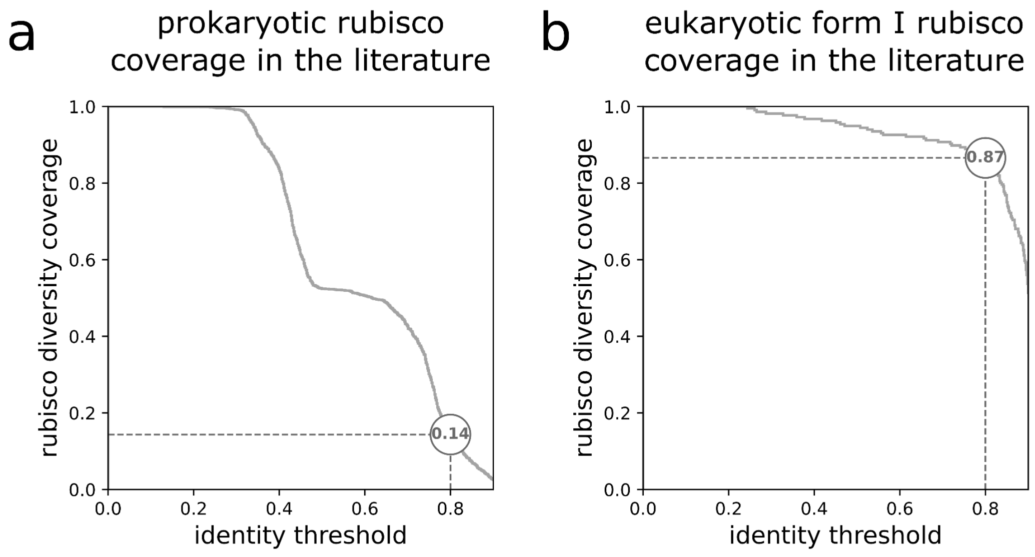


***Fig. S1. Prokaryotic rubisco diversity is poorly covered by the literature in comparison to eukaryotic form I rubiscos, in terms of carboxylation rates.*** *Fraction of rubisco diversity covered at varying sequence identity thresholds in carboxylation rate studies, for (A) prokaryotic and (B) form I eukaryotic variants. Coverage reached at 80% sequence identity is indicated in both panels as an example. Coverage at a given identity threshold, X, is defined as the fraction of representative sequences from a 90% sequence identity clustering that share at least X% identity with any kinetically characterized rubisco variant.*

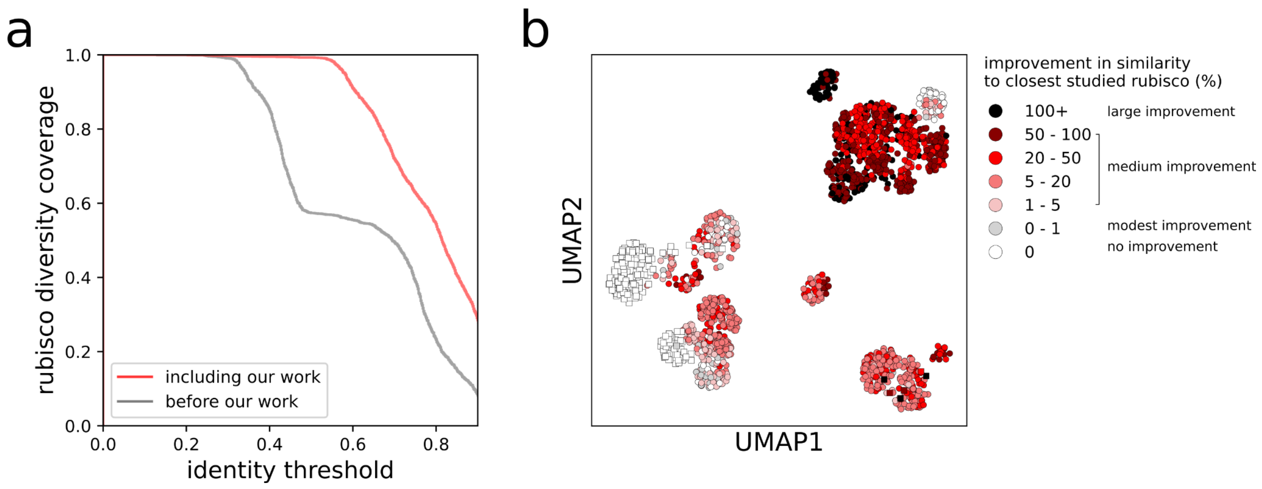


***Fig. S2. Rubisco variants selected in our work dramatically increase rubisco diversity coverage in terms of carboxylation rate study.*** *(A) Fraction of rubisco diversity covered in carboxylation rate studies before and after including variants selected in our work. Coverage at a given identity threshold, X, is defined as the fraction of representative sequences from a 90% sequence identity clustering that share at least X% identity with any kinetically characterized rubisco variant. (B) Improvement in identity to the closest studied variant considering rubiscos selected in our work, for each variant plotted in the UMAP plot as introduced in Fig. 1. Improvement is calculated as the percent identity of each variant to the closest variant from our work minus the identity to the closest variant from previous published works, divided by the latter.*

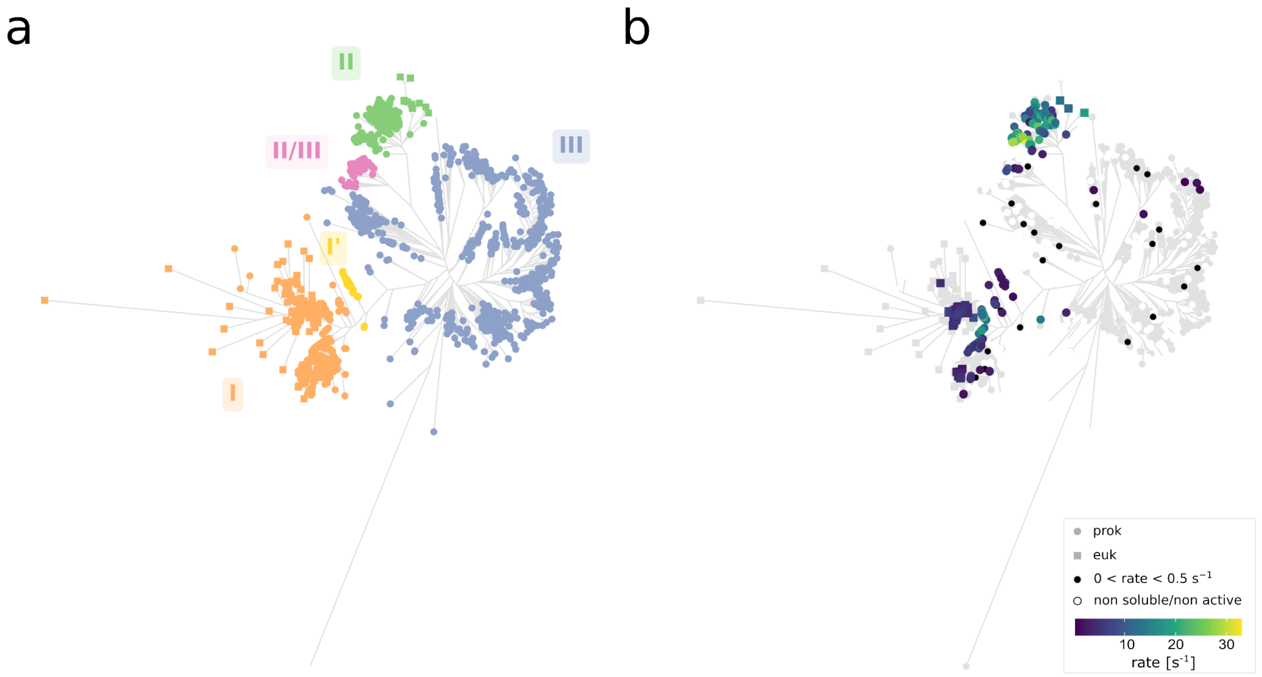


***Fig. S3. Phylogenetic tree presentation of the systematic exploration of rubisco carboxylation rate across its natural genetic diversity.*** *Leaf colors indicate rubisco form (A) or the measured carboxylation rate reported in the literature and/or in our work (B). Prokaryotic and eukaryotic variants are shown as circles and squares respectively. Rubisco variants that were either insoluble or inactive are represented as white circles/squares. Variants with carboxylation rates below 0.5* $s^{-1}$ *are represented as black dots. All rates presented correspond to measurements or corrections made at 30°C.*

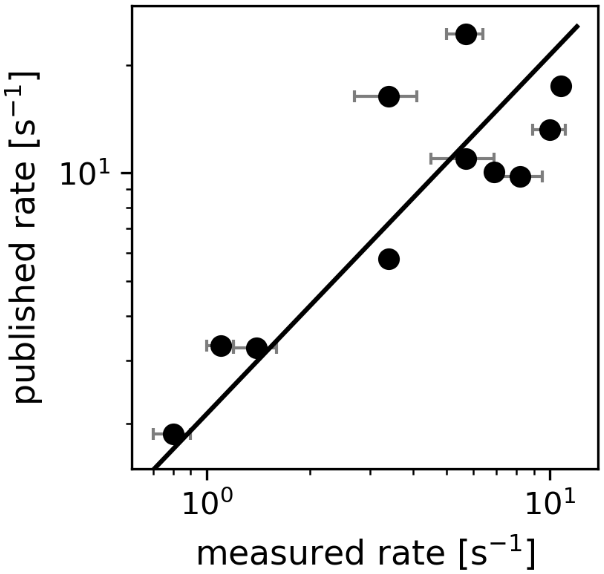


***Fig. S4. Correlation between published and measured carboxylation rates.*** *Measured rates were obtained at 30°C, while published rates were corrected to 30°C. Carboxylation rates measured with a coupled assay tend to be underestimated, compared to direct assays published in the literature. A log–log comparison with a fixed slope of 1 was therefore used to estimate a multiplicative bias (2.1) between our measurements and literature values, which was then applied to correct our measured rates accordingly. Error bars are the standard deviations across our measured rates.* $R^2$ *= 0.68. Pearson correlation coefficient = 0.86.*

***Table S1. Rubisco variants measured in both the literature and in our work.*** *Detail of the values displayed in Fig. S4. The three rates marked with an asterisk (*) were multiplied by two following the recent update published by Tsai et al., 2025 (1).*

| identifier | species | form | rate our lab (30°C) | rate literature (25°C) | median rate literature (corrected to 30°C) | reference |
|---|---|---|---|---|---|---|
| WP_011390153.1 | *Rhodospirillum rubrum* | 2 | 6.9 | 12.3/7.3/5.9/6.28 | 10.1 | (2–5) |
| WP_011500311.1 | *Methanococcoides burtonii* | 2/3 | 0.8 | 1.93/0.6 | 1.9 | (6, 7) |
| WP_011057346.1 | *Thermosynechococcus elongatus* | 1 | 5.7 | 7.4 | 11.0 | (8) |
| WP_012536704.1 | *Acidithiobacillus ferrooxidans* | 1 | 3.4 | 11* | 16.3 | (9) |
| WP_011130576.1 | *Prochlorococcus marinus* | 1 | 8.2 | 6.58 | 9.8 | (10) |
| RME08239.1 | *Anaerolineae bacterium* | 1 | 1.4 | 2.2* | 3.3 | (11) |
| WP_011242444.1 | *Synechococcus PCC6301/PCC 7942* | 1 | 10.8 | 14.4/11.8/11.4/9.78/ 14.4/6.6/6.1/11.8/ 12.1/12.6/9.0/14.3 | 17.5 | (8, 10, 12–21) |
| RMG64267.1 | *Calditrichaeota bacterium* | 1 | 3.4 | 3.88* | 5.8 | (11) |
| WP_012823801.1 | *Halothiobacillus neapolitanus* | 1 | 10.0 | 8.9 | 13.2 | (22) |
| AAC38280.1 | *Endosymbiont of Riftia pachyptila* | 2 | 5.7 | 16.4 | 24.3 | (23) |

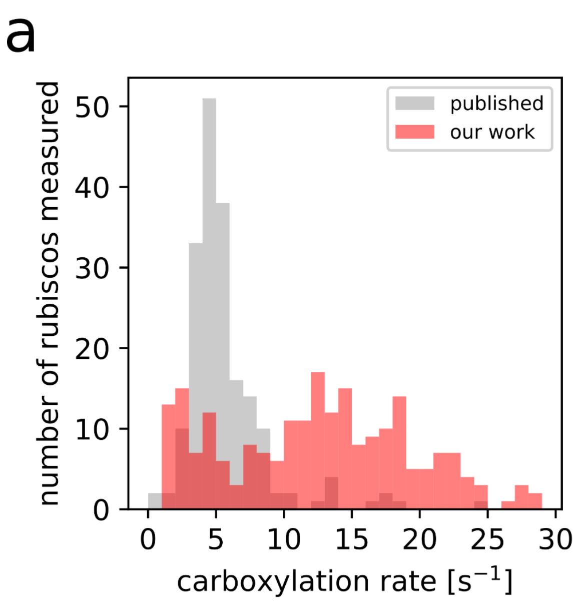


***Fig. S5. Carboxylation rates measured in our work complement previously published data.*** *(A) Histogram of rubisco carboxylation rates measured in the literature (mostly eukaryotic form I) and in our work. (B) Merged datasets show relatively low rates across all forms.*

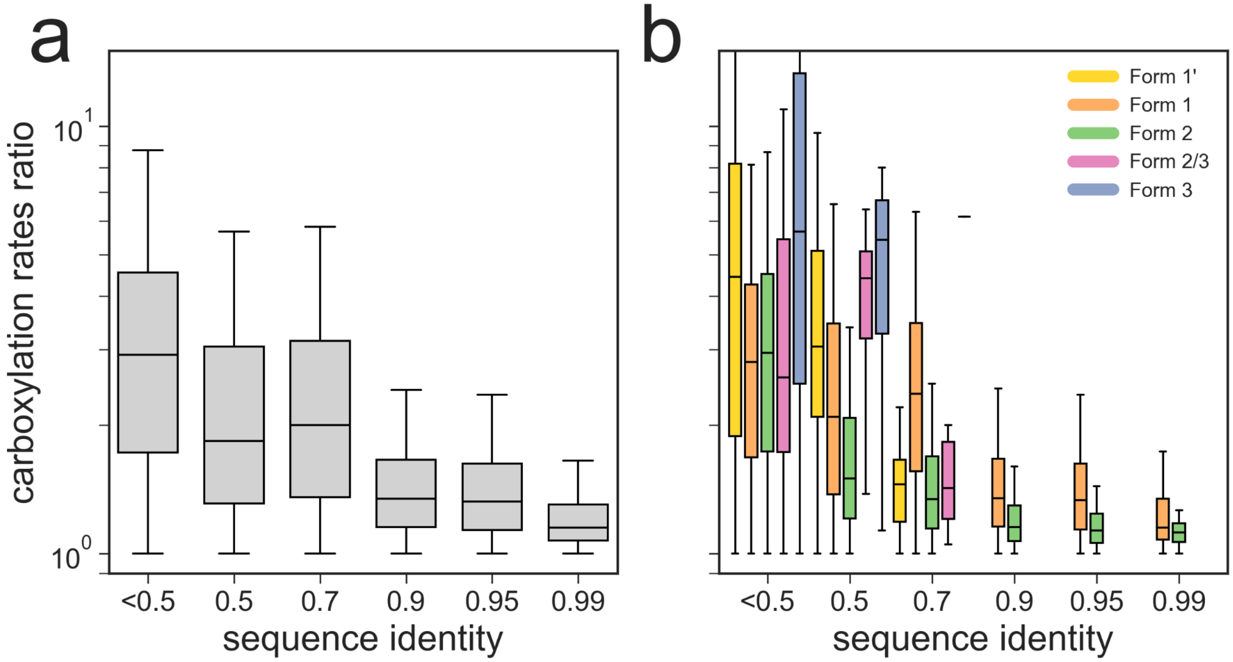


***Fig. S6. Correlation between sequence identity and carboxylation rate ratio among rubisco variants.*** *(A) All pairs combined. (B) Separated by rubisco forms.*

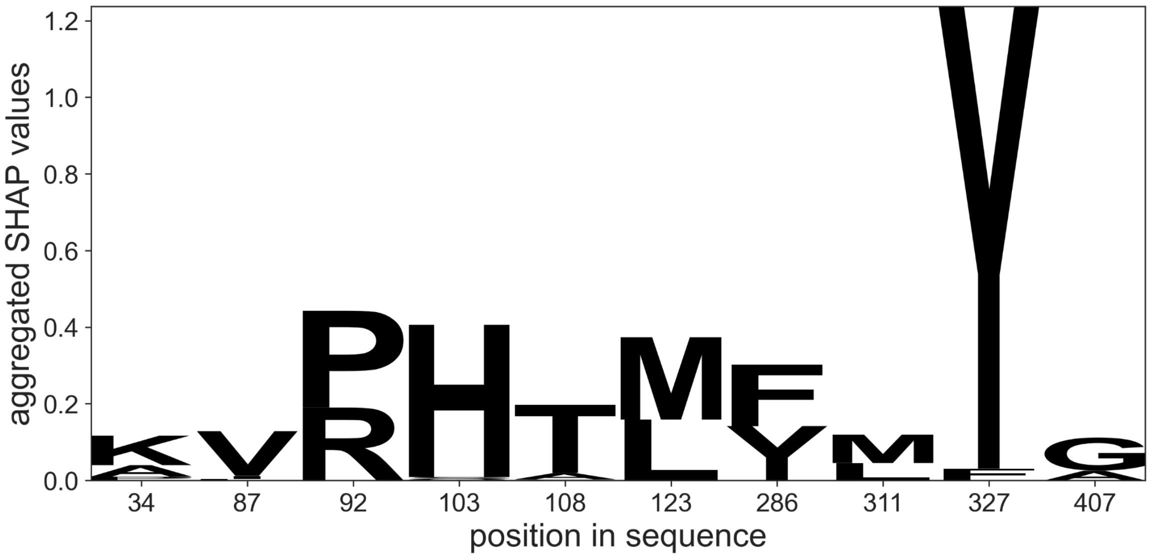


***Fig. S7. Sequence logo representing the 10 main amino acids associated with higher carboxylation rate in rubisco sequence.*** *Random forest regression on the 148 characterized form II and II/III rubisco variants identified the 10 positions with the highest SHAP values, which are represented in this logo, showing the amino acids most strongly correlated with enhanced carboxylation efficiency.*

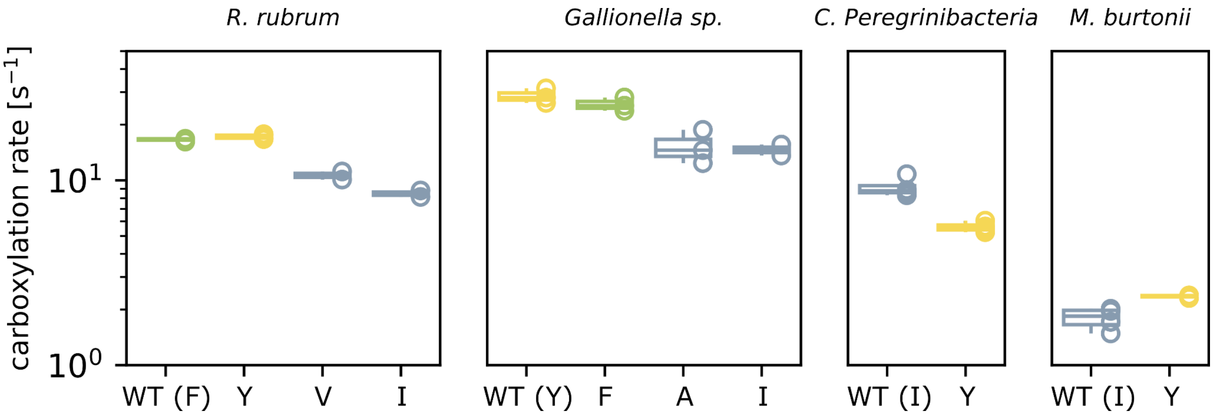


***Fig. S8. Engineering of residue 327 alone does not enhance rubisco carboxylation rate.*** *Measured carboxylation rates of mutants, at residue 327, of four form II rubiscos, from R. rubrum, Gallionella sp., Candidatus Peregrinibacteria, and Methanococcoides burtonii.*

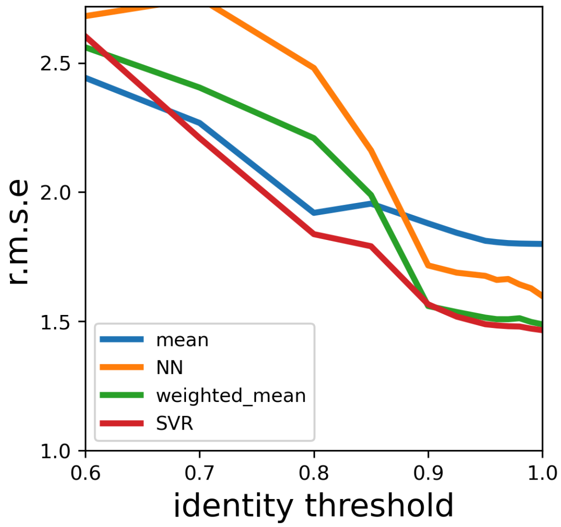


***Fig. S9. Leave-one-out cross-validation of rubisco carboxylation rate predictions using machine learning models.*** *Root mean square error (r.m.s.e., calculated as the exponential of the root mean squared difference between predicted and measured ln($k_{cat,C}$) values) of carboxylation rate predictions for four models—mean (blue line), nearest neighbor (NN - orange line), identity-weighted mean (green line), and support vector regression (SVR - red line)—plotted against the sequence identity shared between the predicted variant and the subset of characterized variants used for training.*

***Table S2. Summary of rubisco forms characteristics and predicted carboxylation rates.*** *Indicated carboxylation rates are the median predicted rates for each form, at 30°C, using a support vector regression (SVR) model. *Values correspond to those obtained by considering only complete sequences. Values in italics, within brackets, indicate those obtained by including all sequences (both complete and partial).*

| form | subunit composition | number of sequences* | median $k_{cat,C}$ [$s^{-1}$]* | phylogenetic distribution |
|---|---|---|---|---|
| I (prok) | $L_8S_8$ | 3456 *[3971]* | 4.4 *[3.6]* | Pseudomonadota, Cyanobacteriota, Chloroflexota, Actinomycetota, Bacillota, Verrucomicrobiota |
| I (euk) | $L_8S_8$ | 7755 *[61054]* | 4.5 *[4.4]* | Viridiplantae, Euglenozoa, Stramenopiles, Rhodophyta, Haptophyta |
| I' | $L_8$ | 52 *[71]* | 2.1 *[2.1]* | Chloroflexota |
| II (prok) | $L_2$ and $L_n$ | 756 *[817]* | 14.9 *[14.9]* | Pseudomonadota, Actinomycetota, Acidobacteriota, Spirochaetota, Bacillota, Verrucomicrobiota |
| II (euk) | $L_2$ and $L_n$ | 7 *[54]* | 12.2 *[12.9]* | Dinoflagellate algae |
| II/III | $L_{10}$ | 137 *[152]* | 3.2 *[3.2]* | Archaea, Candidate Phyla Radiation bacteria |
| III | $L_2$ and $L_n$ | 1386 *[1583]* | 1.9 *[1.9]* | Archaea, diverse bacteria phyla (CPR, Pseudomonadota, Chloroflexota, Bacillota, Acidobacteriota, Nitrospirota, Thermodesulfobiota, Spirochaetota, etc.) |

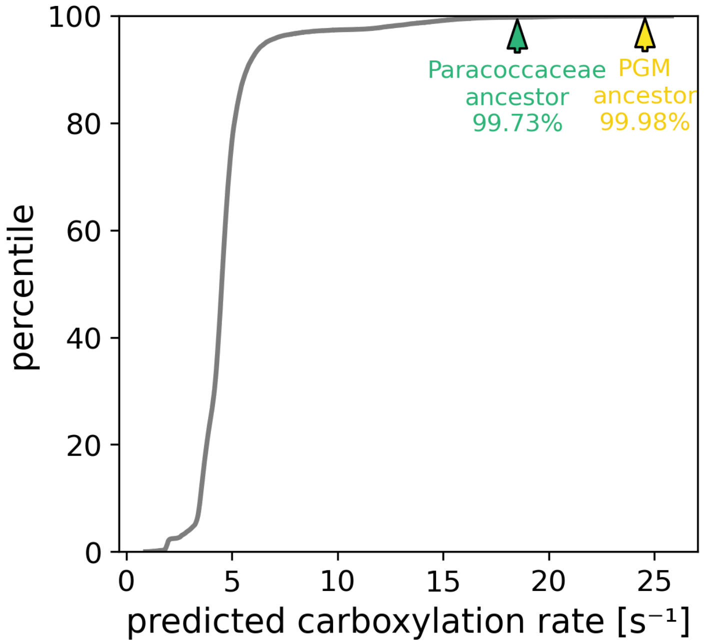


***Fig. S10. Cumulative distribution of predicted carboxylation rates for unannotated rubiscos (grey), showing the percentile positions of the reconstructed Paracoccaceae ancestor (green) and PGM ancestor (yellow) as predicted in a leave one out cross validation of these two variants.***
